# A Systems Approach Identifies Enhancer of Zeste Homolog 2 (EZH2) as a Protective Factor in Epilepsy

**DOI:** 10.1101/836387

**Authors:** Nadia Khan, Barry Schoenike, Trina Basu, Heidi Grabenstatter, Genesis Rodriguez, Caleb Sindic, Margaret Johnson, Eli Wallace, Rama Maganti, Raymond Dingledine, Avtar Roopra

## Abstract

Complex neurological conditions can give rise to large scale transcriptomic changes that drive disease progression. It is likely that alterations in one or a few transcription factors or cofactors underlie these transcriptomic alterations. Identifying the driving transcription factors/cofactors is a non-trivial problem and a limiting step in the understanding of neurological disorders. Epilepsy has a prevalence of 1% and is the fourth most common neurological disorder. While a number of anti-seizure drugs exist to treat seizures symptomatically, none is curative or preventive. This reflects a lack of understanding of disease progression. We used a novel systems approach to mine transcriptome profiles of rodent and human epileptic brain samples to identify regulators of transcriptional networks in the epileptic brain. We find that Enhancer of Zeste Homolog 2 (EZH2) regulates differentially expressed genes in epilepsy across multiple rodent models of acquired epilepsy. EZH2 undergoes a prolonged upregulation in the epileptic brain. A transient inhibition of EZH2 immediately after seizure induction robustly increases spontaneous seizure burden weeks later. Thus, EZH2 upregulation is a protective response mounted after a seizure. These findings are the first to characterize a role for EZH2 in opposing epileptogenesis and debut a bioinformatic approach to identify nuclear drivers of complex transcriptional changes in disease.

**Author Summary:** Epilepsy is the fourth most common neurological disorder and has been described since the time of Hippocrates. Despite this, no treatments exist to stop epilepsy progression. This is fundamentally due to the complex nature of the disease. Epilepsy is associated with hundreds if not thousands of gene expression changes in the brain that are likely driven by a few key master regulators called transcription factors and cofactors. Finding the aberrantly acting factors is a complex problem that currently lacks a satisfactory solution. We used a novel datamining tool to define key master regulators of gene expression changes across multiple epilepsy models and patient samples. We find that a nuclear enzyme, EZH2, regulates a large number of genes in the rodent and patient epileptic brain and that it’s function is protective. Thus, inhibiting EZH2 greatly exacerbates seizure burden. This is the first report of a novel datamining tool to define drivers of large-scale gene changes and is also the first report of EZH2 induction as an endogenous protective response in the epilepsy.

## Introduction

Epilepsy is the fourth most prevalent neurological disorder after stroke, Alzheimer’s disease and chronic migraine^1^. This disease is characterized by excessive and synchronous firing of neurons in the brain that result in the occurrence of spontaneous and unprovoked seizures. Despite its prevalence, the mechanisms governing the manifestation of epilepsy as well as the processes that drive disease progression are poorly understood. A third of all epilepsy patients are refractory to current anti-seizure drugs (ASDs) and prophylactic administration of ASDs does not mitigate disease appearance or progression^2,3^.

Epileptogenesis is the process that links brain insults or pre-disposing genetic mutation(s) to the subsequent emergence of spontaneous seizures. One of the many triggers of epileptogenesis is Status Epilepticus (SE), defined as a seizure lasting more than 5 minutes or multiple seizures occurring without regaining consciousness ^4-6^. SE is followed by a seizure free period termed the latent period. Changes at the molecular, cellular and network levels during the latent period lead to a persistent reduction in seizure threshold resulting in spontaneous seizures.

A lack of understanding of epileptogenic mechanisms has hindered progress towards treatments that target disease progression. Modulation of metabolism^7^, inflammation^8^, chromatin methylation^9^ and signal transduction pathways^10^ have shown promise but attempts to use transcriptome profiling to uncover molecular changes in epilepsy have had varying degrees of success^11-14^. They tend to underscore the important role of inflammation in epilepsy but no other unifying themes have come to light. This is likely due to a combination of factors including differing models, lab practices and low statistical power. Further, with few exceptions^15^ there has been little attempt to define the coordinating factors behind the large-scale gene changes observed in transcriptomic studies.

Transcription factors and their associated cofactors co-ordinate the regulation of hundreds or thousands of genes and it is likely that the majority of gene changes in epilepsy transcriptomic analyses could be explained by the altered function of a small handful of nuclear proteins. This opens up the possibility of normalizing gene expression in disease by targeting a single transcription factor or cofactor, many of which are enzymes, rather than chasing multiple individual differentially expressed genes.

In order to discern those transcription factors and cofactors that drive large scale gene changes during the early latent period we have made use of a recent study that collected transcriptomic profiles from laser captured dentate granule cells in 3 different rodent epilepsy models across 11 laboratories at 3 time points in the early latent period^16^. We have used genome wide chromatin binding profiles (ChIPSeq) of transcription factors and cofactors to screen for nuclear proteins that coordinate gene expression in the latent period. Using a systems approach to integrate ChIPseq and transcriptome profiles we identify the histone methylase Enhancer of Zeste Homolog 2 (EZH2) as a master transcriptional regulator during epileptogenesis. We show that EZH2 levels are significantly increased after Kainic Acid (KA) induced SE in rodent preclinical models and that inhibiting its function worsens seizure burden in mice. We provide evidence that EZH2 function is elevated in human TLE. This is the first study to identify a role for EZH2 in controlling epileptogenesis and uncovers an innate protective mechanism in epilepsy.

## Results

### Status Epilepticus instigates an enduring transcriptional program in dentate granule cells

We utilized the recently published transcriptome profiles of dentate granule cells in epileptic rats (GSE47752) to define transcriptional changes that occur during the early latent period^16^. This is a well-powered expression dataset provided by the Epilepsy Microarray Consortium, who examined the transcriptional profile of laser captured dentate granule cells 1, 3, and 10 days after SE in 3 rat models of SE. Data from the pilocarpine and kainate models were each provided by two independent laboratories, and a single laboratory provided data from the self sustaining status epilepticus model. For each time point, we extracted the median transcript level from each laboratory and model for every gene and filtered for genes expressed above background (see Methods). Fold changes and probabilities were assigned for each expressed gene and differentially expressed genes were defined as those with a fold change at least 2 Standard Deviations greater than the mean fold change *plus* an associated FDR<0.05. Across the 3 models, we found that of the 9614 genes expressed in dentate granule cells, 482 genes were differentially expressed under the above criteria at 1d post SE (243 induced, 239 repressed) (**Figure 1A**). 282 genes were induced and 170 repressed at d3 (**Figure 1B**) and 235 were induced and 223 repressed at 10d (**Figure 1C**) (**Supplemental Table 1**). As seen in Figures 1A-1C, differentially expressed genes so defined, were able to segregate control from epileptic samples at all 3 time points.

**Figure 1:**
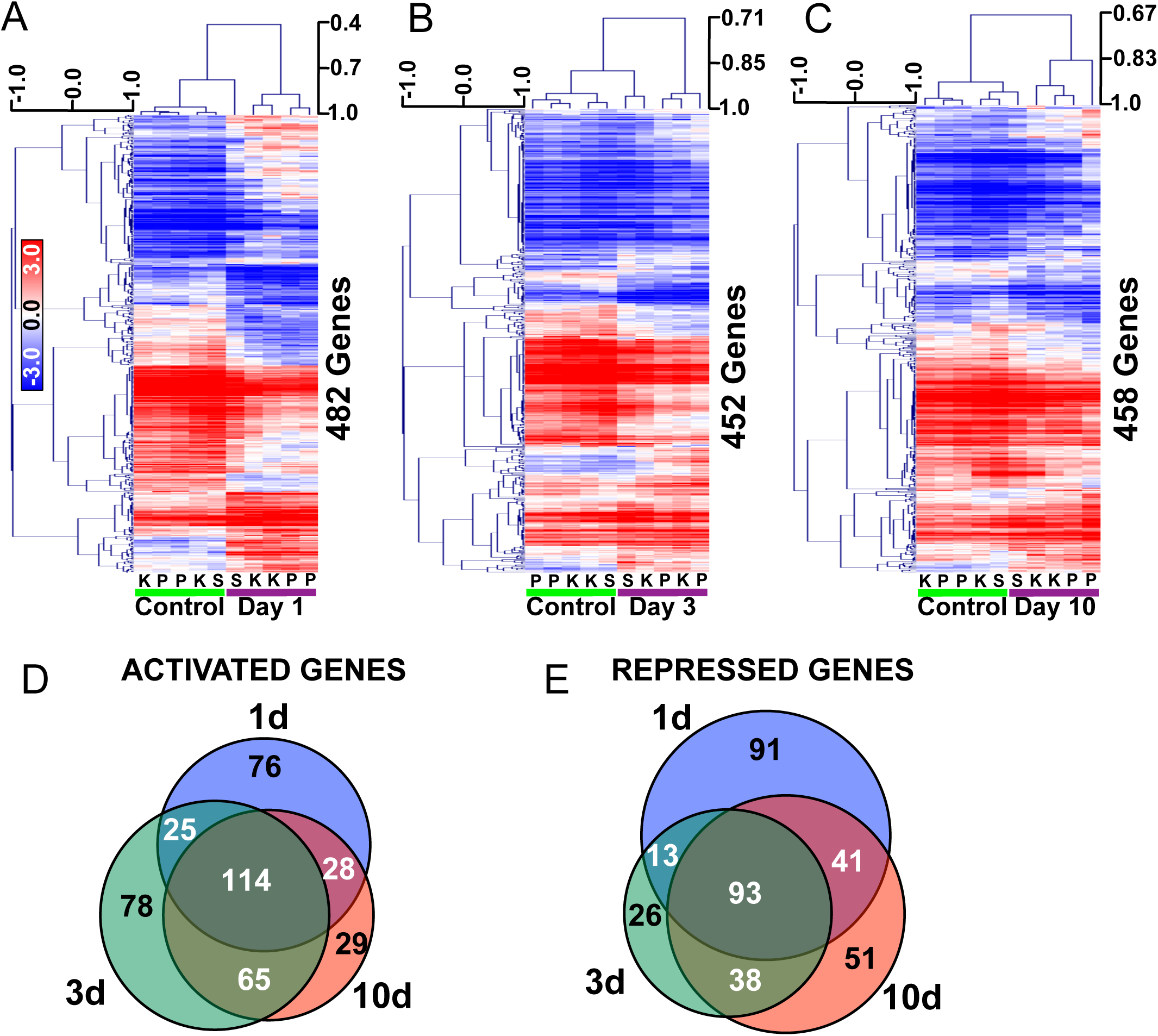
Transcriptome analysis of rat dentate granule cells post SE highlights persistent gene changes in the early latent period. A-C) Heat maps of genes differentially expressed by more than 2 Standard Deviations of the mean change and FDR<0.05 at 1,3, or 10d after SE induced by systemic Kainate (k), Pilocarpine (p) or Self-Sustaining Status Epilepticus (s). Scale: Pearson correlation coefficient. D) Venn diagram showing overlap between genes up-regulated 1d, 3d and 10d post SE. E) Venn diagram showing overlap between genes down-regulated 1d, 3d and 10d post SE.

Interestingly, of the 243 genes that were induced at 1d, 142 remained induced at day 10 (p=7.3×10^−183^, odds ratio=139, 58% of 1d and 60% of 10d genes). Of the 282 genes that were elevated in day 3, 179 remained so by day 10 (p=1.2×10^−251^, odds ratio=283, 63% of 3d and 76% of 10d genes) (**Figure 1D**). Similarly, of the 239 repressed genes at 1d, 134 remained repressed 10 days after SE (p=2.2×10^−172^, odds ratio=133, 56% of 1d and 60% of 10d) and of the 170 genes repressed at 3d, 131 remained repressed 10 days after SE (p=2.4×10^−197^, odds ratio=341, 77% of 3d and 59% of 10d) (**Figure 1E**).

Ontological analysis on the 142 persistently upregulated genes suggests an ongoing inflammatory response. Genes associated with terms such as ‘signaling by interleukins’ and ‘cytokine signaling in Immune systems’ were enriched in the activated gene list (**Supplemental Table 2**). The 134 persistently repressed genes were not as significantly enriched in any one GO term (FDR > 5%) but were somewhat associated with cell-cell communication with terms such as ‘Adherens junctions interactions’ and ‘cell junction organization’ predominating (**Supplemental Table 3**).

### EZH2 is a driver of transcriptional changes in the early latent period

A single transcription factor or cofactor (we shall refer to both as “Factors” herein) can regulate many hundreds or thousands of genes^17^. We therefore hypothesized that the hundreds of differentially expressed genes post SE may be regulated by one, or a few, Factors. To test this, we made use of genome wide Factor binding data archived at the Encyclopedia of DNA Elements^18^ that consists of 161 ChIPseq tracks for 119 Factors across 91 cell types. We first assigned a ChIP signal for every Factor to every gene in ENCODE by determining the highest ChIP signal found for that Factor at every locus across all cell type resulting in a matrix of ChIP values for each Factor at every gene (see Methods). We then identified those Factors that were preferentially ChIPed at our differentially expressed gene lists. This was accomplished by assigning a score based on the likelihood that the differentially expressed genes were biased towards high ChIP signals and how the highest ChIP signals in the differentially expressed genes compare to the highest signals across all 9614 expressed genes. Factor analysis of persistently up-regulated genes highlighted 11 Factors as having significantly higher ChIP signals in the input list compared to the 9614 expressed genes (FDR<5%) (**Figure 2A, Supplemental Table 4)**. The highest scoring Factor was STAT3 (FDR=3×10^−5^, Score=3.41) which is consistent with the ontological analysis showing an enrichment for inflammatory genes^19^. Analysis of persistently down-regulated genes highlighted 9 Factors with EZH2 being the highest scoring (FDR=2.8×10^−24^, Score=12.4) (**Figure 2B, Supplemental Table 5**).

**Figure 2:**
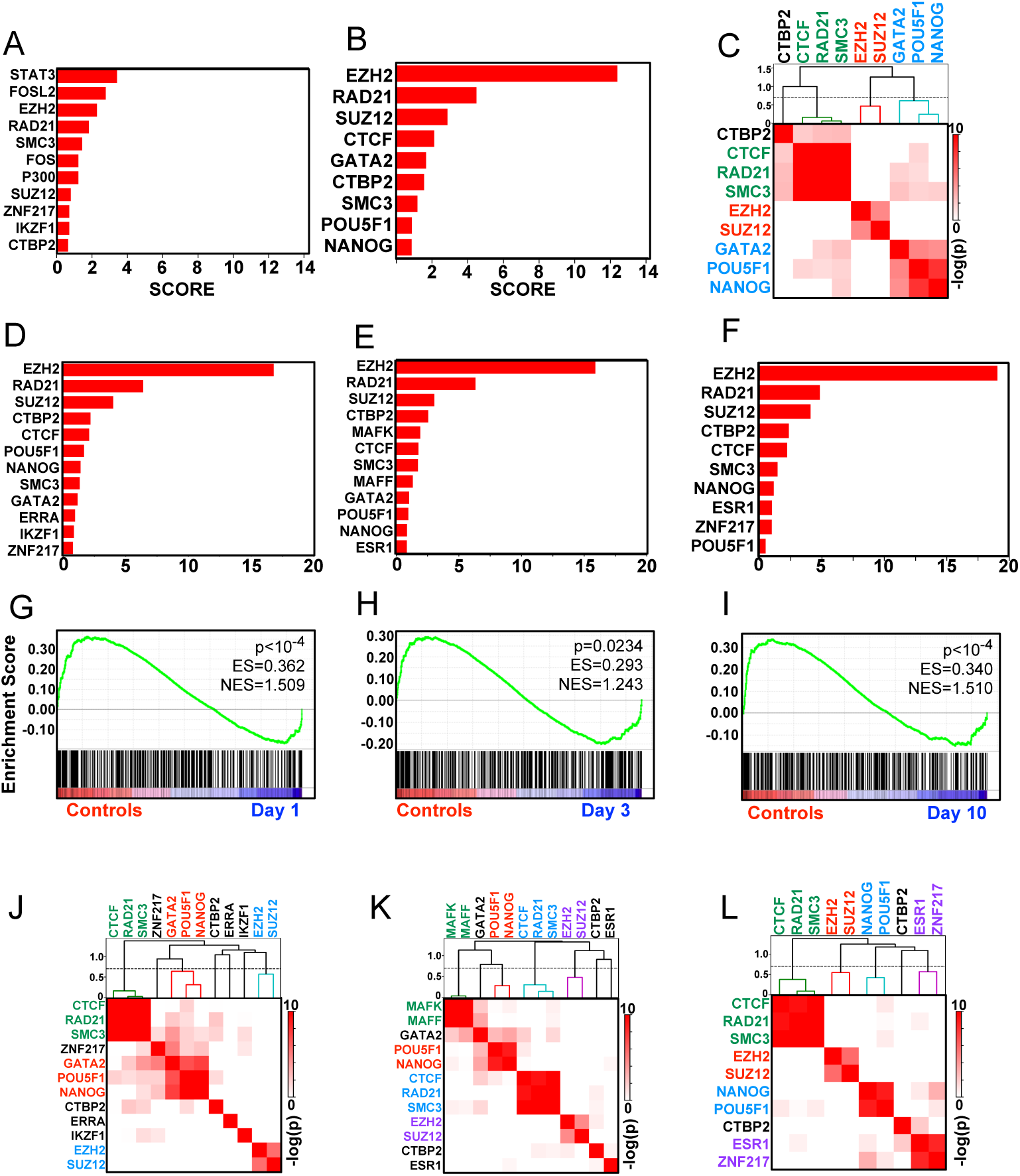
EZH2 drives gene change repression in the early latent period. A) Genes upregulated at both days 1 and 10 are enriched for STAT3 ChIP peaks across ENCODE ChIP-seq profiles. B) Persistently repressed genes are enriched for EZH2 peaks across ENCODE ChIP-seq profiles. C) Overlaps between lists of target genes for each significant Factor are plotted as a heatmap based on Fisher Exact test p values. Clustering by Pearson distance highlights 3 groups of factors controlling persistently repressed genes. Dashed line cutting branches of the hierarchy tree at Pearson distance=0.7 (Pearson correlation=0.3). D-F) Factor analysis highlights EZH2 as the principle driver of repression 1d, 3d or 10d post SE. G-I) GSEA shows EZH2 target genes are enriched in control samples and depleted in repressed genes 1d, 3d or 10d post SE. J-L) Fisher Exact tests of driven genes 1d, 3d or 10d post SE shows EZH2 and SUZ12 co-target genes at all 3 timepoints.

Given the robust EZH2 FDR and Score compared to STAT3, and its established role in epigenetic silencing in other systems^20-22^, we further explored the regulation of persistently repressed genes by EZH2. We asked whether the 9 Factors (EZH2, RAD21, SUZ12, CTCF, GATA2, CTBP2, SMC3, POU5F1, NANOG) that target repressed genes all target the same genes, or whether each Factor targets a unique subset of persistently repressed genes. For each Factor we defined target genes using an approach analogous to the ‘Leading Edge’ analysis in Gene Set Enrichment Analysis (GSEA)^23,24^. Thus, target genes were defined as that subset of repressed genes whose ChIP signals were greater than the argument of the Kolmogorov-Smirnov statistic for the cumulative density functions for all background genes (i.e. the 9614 expressed genes) and all repressed genes (see Methods and **Supplemental fig. 2**). Hypergeometric probabilities were assigned to all pairwise comparisons of Factor target genes with an odds ratio>1. This resulted in a 2-dimensional array of probabilities that any 2 Factors’ target genes have a significant overlap. Factors were then clustered based on likelihoods of sharing target genes. **Figure 2C** shows that the Factors targeting persistently repressed genes group into distinct clusters. EZH2 and SUZ12 have a significant overlap of target genes (p=2×10^−5^, odds ratio=100, Fischer Exact). This is consistent with the pair acting as part of the Polycomb Repressive Complex 2 (PRC2)^25,26^. CTCF, SMC3, and RAD21 also co-target genes and would reflect their function as the core of the Cohesin complex^27^. NANOG, GATA2 and POU5F1 also form a cluster consistent with their known role in stem cell regulation^28^. Performing Factor analysis on genes repressed on day 1, day 3 or day 10 post SE showed that EZH2 targets were the most enriched in all 3 days (**Figure 2D-F, supplemental tables 6-8**).

To further test the hypothesis that EZH2 targets are enriched in the persistently repressed genes, we generated a gene set consisting of the most highly ChIPed EZH2 genes in ENCODE (see Methods) and then performed GSEA using the transcriptomes from 1,3 and 10 days post SE. **Figure 2G-I** show that EZH2 targets are enriched in control samples and under-represented at the 3 time points post SE. At all 3 time points, EZH2 target genes had a significant overlap with SUZ12 targets (**Figure 2J-L**). These analyses support the hypothesis that genes bound by EZH2 are repressed during the early latent period and that EZH2 likely functions in concert with SUZ12 to target repression after SE.

### EZH2 regulates a gene co-expression network in Human Temporal Lobe Epilepsy

To determine whether EZH2 might coordinate gene expression in human epilepsy, we made use of whole transcriptome data from 129 Temporal Lobe Epilepsy (TLE) ante-mortem samples first described by Johnson et al^15^. We used k-medians clustering to generate 10 gene clusters (modules M-1 to M-10), each containing co-expressed genes across the 129 samples. Each gene cluster was then tested to see if any had genes that were preferentially bound by EZH2 using the Factor analysis algorithm. EZH2 was the top scoring factor for cluster M-1 (Score=15.2, FDR=4.6×10^−20^). Indeed, only M-5 had a higher scoring factor than EZH2 in M-1 (TAF1; Score=16.0, FDR=2.8×10^−19^) (**Figure 3A**). The other factors preferentially binding M-1 (**Figure 3B, Supplemental Table 9**) showed significant overlap with those factors driving down-regulation of genes in rats in the early stages of epileptogenesis (**Figure 3C**). Thus, when we performed unsupervised clustering of TLE gene clusters and genes up- or down-regulated in epileptic rats based on the factors controlling those genes, M-1 segregated with rat gene lists that were down-regulated at days 1,3, and 10 post SE (**Figure 3D and Supplemental Figure 1**).

**Figure 3:**
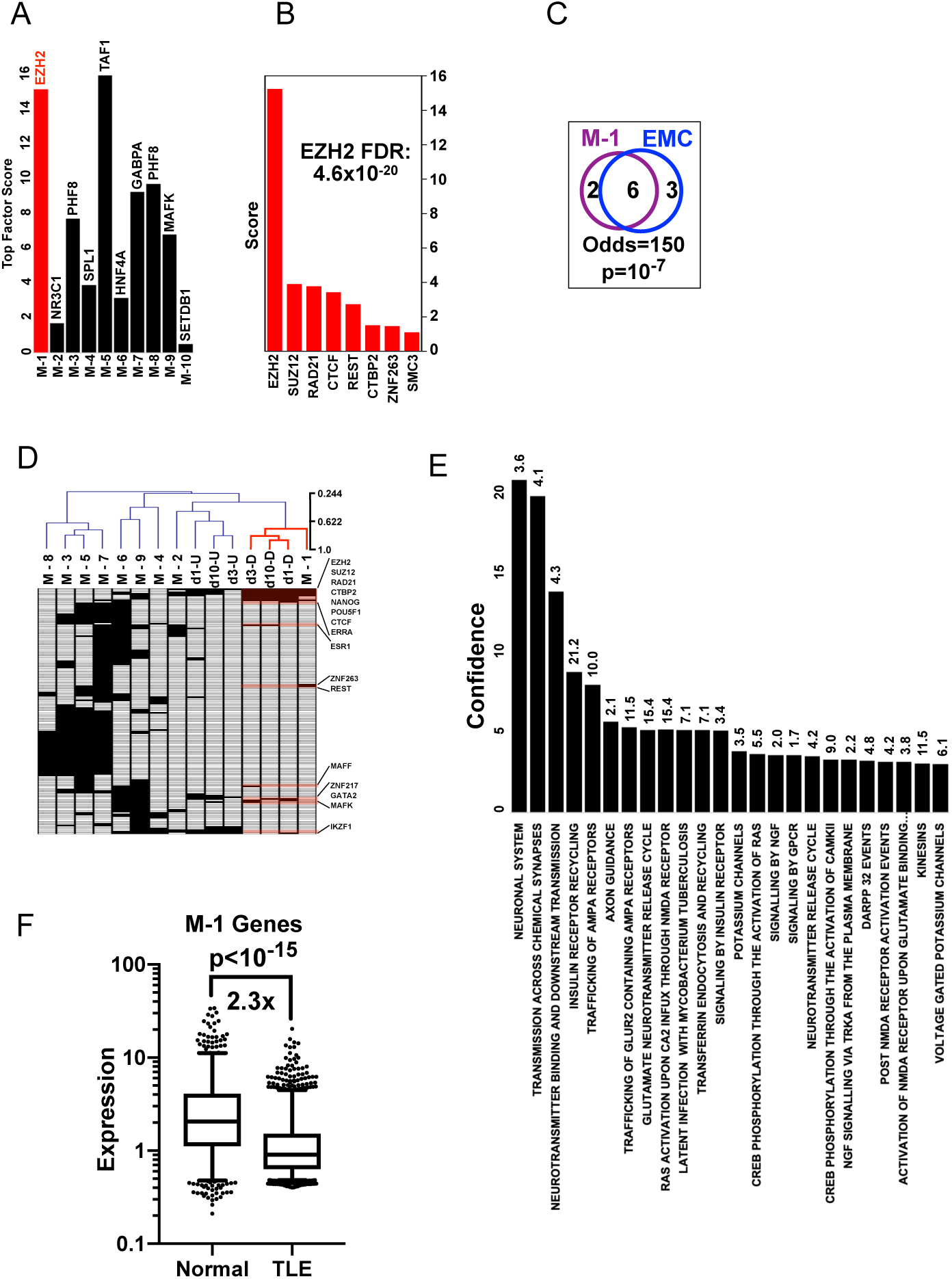
EZH2 targets a module of genes in human TLE. A) Top scoring Factors are plotted for the 10 k-median clusters in human TLE. B) All positive Factors for module M-1. C) Venn diagram of Factors positive for M-1 and positive for down-regulated genes on day-1 in the rodent EMC data with Fisher Exact p value and odds ratio. D) All 10 k-median human TLE modules and rat gene lists for up or down regulated genes on days 1,3 and 10 after SE were clustered based on absence or presence of factors in their respective Factor analyses – only those factors positive for human M-1 or rat days 1,3 and 10 are shown. The complete map with all factors is shown in supplemental figure 1. Black box denotes Factor with FDR<0.05, white denotes Factor with FDR≥0.05. E) Ontological analysis of M-1 using Reactome database (odds ratios shown above each column). F) Expression of M-1 genes in non-epileptic (‘Normal’) versus TLE human samples.

Ontological analysis using the Reactome database^29^ shows that M-1 genes are enriched for terms associated with neuronal transmission, signaling, receptor trafficking, and neurotransmitter release (**Figure 3E**).

Thus, EZH2 emerges as a factor associated with seizures in both humans and rat. To begin testing the function of EZH2 in human TLE, we compared the expression levels of M-1 genes in the above TLE samples and 55 post mortem hippocampal transcriptomes from individuals with no psychiatric or neurological disorders, substance abuse, or any first-degree relative with a psychiatric disorder as described in Li et al (GSE45642)^30^. Consistent with an increase in EZH2 corepressor function, M-1 genes had lower expression in TLE versus non-epileptic samples (**Figure 3F**). Johnson et al identified a group of 442 genes in the TLE dataset using Graphic Gaussian Modelling^15^. The group was enriched for genes associated with inflammation and immune response. We found that cluster M-4 was highly enriched for genes controlling cytokine interactions and immunity (**Supplemental Table 10**). Further, our Factor analysis highlighted a number of factors known to control inflammation including IKZF1 and NfKB (**Supplemental Figure 3 and Supplemental Table 11**). Consistent with the pathological role of inflammation in epilepsy, we found that the normal hippocampus did not yield any group of co-regulated genes that overlapped with M-4 or that were enriched for terms including ‘immunity’ or ‘inflammation’ or driven by known factors controlling inflammation such as IKZF1 or NfKB.

### EZH2 protein levels are increased after SE

To test whether the increased EZH2 function predicted by our Factor analysis corresponds to increased EZH2 expression *in vivo*, we utilized the low-dose systemic KA mouse model. Hippocampi were analyzed from saline injected and KA induced mice at eight different time points (4 hours, 1 day, 2 days, 4 days, 5 days, 10 days, 20 days, and 30 days) after SE (**Figure 4A**). These time points were chosen to capture a representative profile of the acute to more chronic changes that occur after SE^31^. Western blot of whole hippocampus shows a robust and prolonged increase in EZH2 protein 2 to 5 days after SE, peaking at 6-fold above saline injected controls on Day 2 (**Figure 4B**, quantified in **Figure 4C**). EZH2 protein trended back to saline control levels by 10 days.

**Figure 4:**
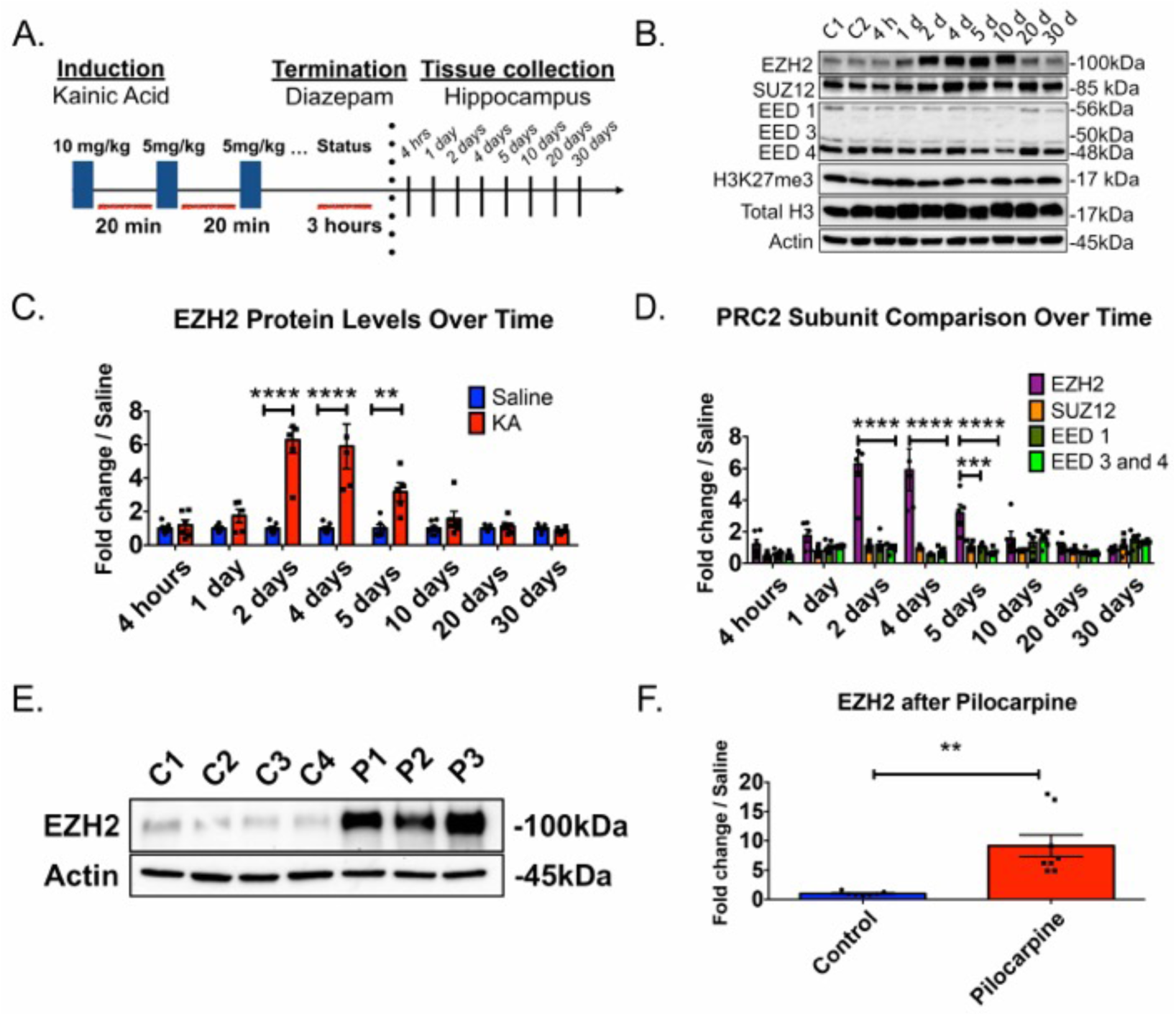
EZH2 protein levels are increased 2-5 days after KA induced SE. (A) Schematic of low dose KA protocol used for Figures 4-7. See Methods for complete description. (B) Representative Western Blot of EZH2, SUZ12, EED-1, EED-3, EED-4, H3K27me3 and Total H3 levels after SE, where C1 = untreated animals, C2= saline treated animal. Lanes 3-10 represent different KA treated mice sacrificed at time points post SE. C = control, H = hours, D = days. Quantifications of SUZ12, EED-1, EED-3, EED-4, H3K27me3 and Total H3 are located in supplemental information. (C) Quantification of EZH2 protein western blots across time. EZH2 exhibits a robust, prolonged increase in protein levels beginning with a 6.3-fold increase compared to saline animals 2 days after SE. This increase remains statistically significant out to 5 days. (D) Graph demonstrates fold change in EZH2 protein levels relative to other PRC2 subunit components. Relative fold change of EZH2 in KA animals is significantly greater than fold changes detected for SUZ12, EED-1 and EED-3 and EED-4 proteins at 2, 4 and 5 days. (E) Representative Western blot of hippocampi extracted 48 hours after pilocarpine treatment in rats. C = saline treated controls. P = Pilocarpine treated. See methods for complete description of induction paradigm used. Each band represents one hippocampal hemisphere from one rat. We find that EZH2 levels are up regulated 14.7-fold after SE. (F) Quantification of EZH2 protein from all tested pilocarpine rats (n = 7 saline controls, n = 8 pilocarpine). For panels C-D, statistical significance was assessed across animals, time, and conditions through Two-way ANOVA test where p < 0.05 with Holm-Sidak correction for multiple comparisons. n = 7 for saline treated animals and n = 5-7 KA animals at every time point. For Panel F, statistical significance was assessed by unpaired Student’s t-test. *p <0.05, **p < 0.01, ***p< 0.001, ****p<0.0001.

EZH2 requires the presence of its partner components in the Polycomb Repressive Complex II, SUZ12 and Embryonic Ectoderm Development (EED), to catalyze tri-methylation of lysine 27 on histone H3 (H3K27me3)^26,32^. To characterize how levels of these proteins and histone marks change post SE, we performed Western Blot analysis on SUZ12, EED-1, EED-3, EED-4, H3K27me3 and Total H3. We find that SUZ12 protein is transiently downregulated at 4 hours after SE but is similar to controls at other time points tested (**Figure 4B, 4D** and **Supplemental Figure 5A**). EED-1 protein levels showed no change (Figures 4B, 4D and **Supplemental Figure 5B**), while EED-3 and EED-4 showed a transient down-regulation at 4 hours after SE, and an increase at 10 days (**4B, 4D** and **Supplemental Figure 5C**). Levels of H3K27me3 were globally up-regulated at 1 day and 10 days (**Supplemental Figure 5D**). In summary, the robust induction of EZH2 post SE is not mirrored by the other PRC2 subunits.

**Figure 5:**
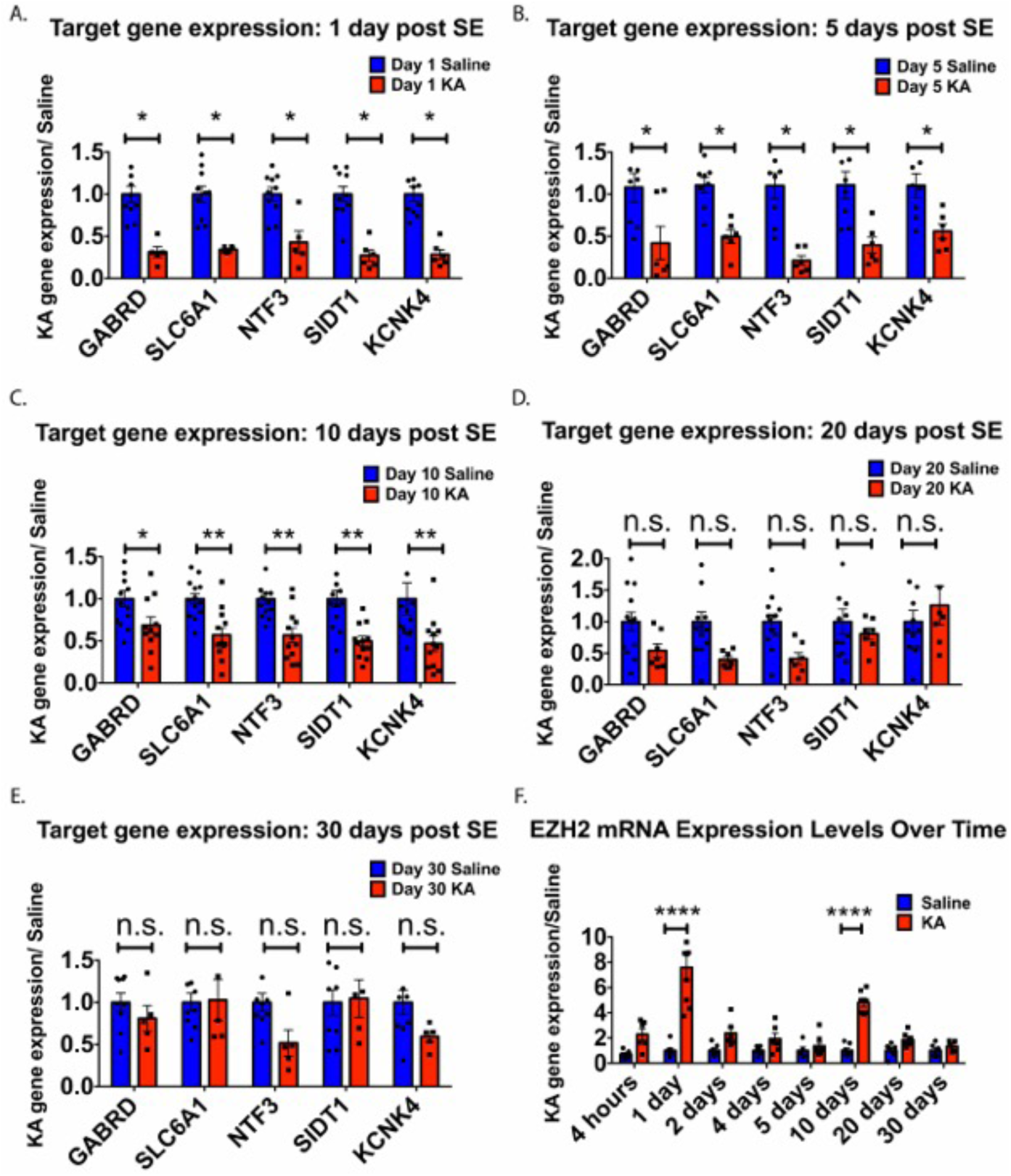
EZH2 mRNA levels exhibit early, transient increases after SE, while EZH2 target gene expression is down regulated out to 10 days. For panels (A-E), gene expression data for five tested EZH2 target genes are shown. All statistical tests were performed using multiple ttests with Holm-Sidak correction for multiple comparisons.*p <0.05, **p < 0.01, ***p< 0.001, ****p <0.0001. n = 3-13 animals for saline and n = 5-13 animals for KA conditions. Target genes show significant down regulation after SE out to 10 days (A-C), with the exception of SLC6A1 and KCNK4 showing no difference compared to saline animals at day 5. Down regulation in SLC6A1 and KCNK4 returns at 10 days (C). By 20 and 30 days, differences in gene expression are abated (D-E). (F) EZH2 mRNA levels significantly increase at 1 day and 10 days post SE. Statistical significance was assessed by Two-way ANOVA and multiple comparisons were corrected for by Holm-Sidak correction. n = 6-7 animals for saline and n = 5-9 animals for KA conditions.

To assess whether EZH2 induction is restricted to the KA mouse model or occurs across models, EZH2 protein was monitored after SE in a lithium-pilocarpine rat model. **Figures 4E** show that EZH2 is also up-regulated in the hippocampus of pilocarpine-treated rats, with a 14.7-fold increase compared to saline injected controls (**quantified in Figure 4F**). Thus, we observe EZH2 is up regulated in two different rodent species (mouse v. rat) and in two different post SE induction paradigms (KA v. pilocarpine).

### EZH2 target genes are down regulated post SE

To test whether the EZH2 induction is functional, we performed Quantitative Reverse Transcription PCR (qRT-PCR) on five EZH2 target genes identified in the Factor analysis (**Supplemental tables 12,13,14**). The chosen target genes have important functions in glutamatergic and GABAergic signaling and have been implicated in epilepsy disease pathology ^33,34^. We found that these EZH2 target genes were down regulated up to 10 days after SE (**Figure 5A-C**). Expression levels begin to increase back to saline levels at 20 days and are fully restored by 30 days (**Figure 5D,E**).

The increase in EZH2 protein in **Figure 3** led us to ask whether increases in EZH2 are a result of a transcriptional or post-transcriptional mechanism. **Figure 5F** shows that EZH2 mRNA levels exhibit an early, transient increase 1 day after SE, returning to control levels between days 1-5 and increased again at 10 days. This result suggests a complex mechanism of EZH2 induction that may involve a transcriptional component.

### Inhibition of EZH2 *in vivo* increases seizure burden in KA mice

The definitive characteristic of epileptogenesis is the generation of spontaneous, recurrent seizures (SRS)^35-38^. To test whether EZH2 up-regulation affects SRS, we intraperitoneally (i.p.) delivered UNC1999 ^39^, a specific pharmacological inhibitor of EZH2, to mice after SE induction for 3 days and monitored seizure progression 5 weeks later. We chose to deliver the drug post SE to avoid potential interference of UNC1999 with the manifestation of SE itself.

FVB/NJ mice were induced as described above and dosed with 20 mg/kg of UNC1999 or drug vehicle, beginning 6 hours post SE induction and then twice more at 24 and 48 hours. The timing of dosing was determined based on the first characterization of UNC1999 by Konze *et al* in 2013, who demonstrated that treatment of UNC1999 for three consecutive days reduces H3K27me3 levels *in vitro*^39^. The dose was determined prior to experimentation using a dose toxicity study on naïve FVB/NJ male mice. Pain and distress were assessed using the NIH guidelines for Pain and Distress in Laboratory Animals: Responsibilities, recognition, and Alleviation (https://oacu.oir.nih.gov/sites/default/files/uploads/arac-guidelines/pain_and_distress.pdf). 20 mg/kg was found to the best tolerated dose by naïve mice.

Five weeks post SE, during which KA treated mice develop spontaneous seizures, mice from each treatment group were video recorded for 8 hours daily, 6 days a week for 3 weeks (**Figure 6A**). Videos were scored by two individuals blinded to treatment groups utilizing a modified version of the Racine scale (see Methods)^40^.

**Figure 6:**
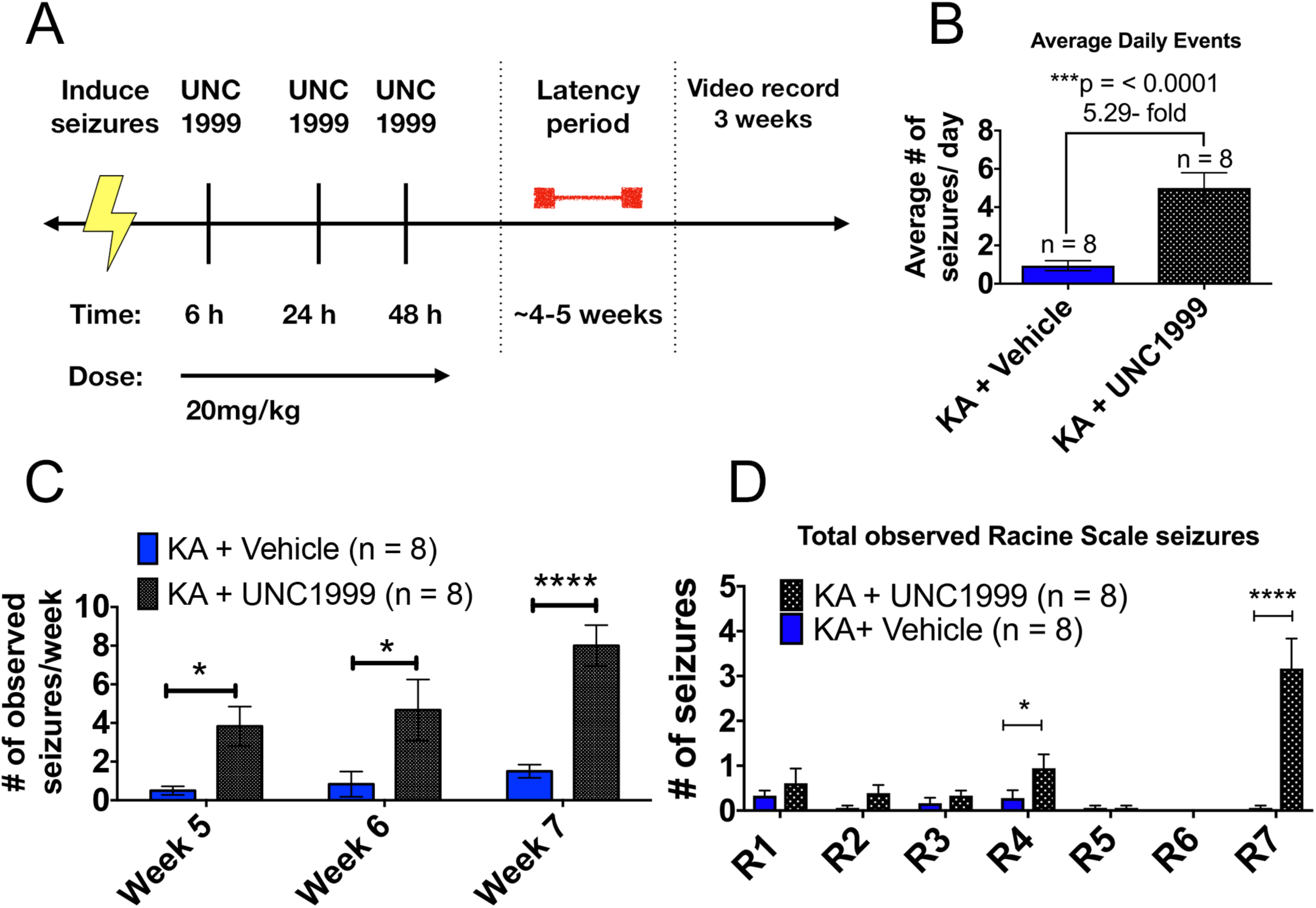
Inhibition of EZH2 *in vivo* increases seizure burden in KA treated mice (A) Schematic of UNC1999 dosing regimen used with KA to test animals. Briefly, animals were induced to SE and then dosed three times with 20mg/kg UNC1999 at 6, 24 and 48 hours. Animals were allowed to recover for 4-5 weeks and then scored through video monitoring for 3 weeks. Two observers blinded to treatment groups scored all videos. (B) KA + UNC1999 exhibited 5.29-fold more average seizures per day compared to vehicle treated animals (Student’s unpaired t-test with equal SD, p <0.05). (C) KA+ UNC1999 treated animals demonstrate significantly greater number of seizure events per week of recording compared to vehicle treated mice (Two-way ANOVA test, p< 0.05, with Holm- Sidak correction). (D) KA + UNC1999 treated mice exhibited significantly more rearing and running, jumping and spinning in circles behavior compared to vehicle, suggesting a more severe disease progression. Statistical significance assessed by Student’s unpaired t-test, n = 6 saline + vehicle, n = 3 KA + vehicle and n = 7 KA + UNC1999 animals. *p <0.05, **p < 0.01, ***p< 0.001, ****p <0.0001.

Transient inhibition of EZH2 via UNC1999 for 3 days post SE significantly increased the number of daily seizure events in KA mice 5 weeks later (**Figure 6B**). UNC1999 treated KA mice also exhibit seizures that are more severe and increase in number as the weeks progressed (**Figure 6C**). The largest increase in Racine scale behaviors displayed by UNC1999 treated animals included rearing (R4) and violent running, jumping or spinning in circles (R7) with several continuous minutes of spinning in a clockwise manner (**Figure 6D**). Administering UNC1999 alone failed to elicit seizures arguing that the drug itself is not epileptogenic (**Supplemental figure 6A**). These data are consistent with the hypothesis that increased EZH2 post SE is a protective response and that antagonizing EZH2 exacerbates epilepsy.

### UNC1999 crosses the BBB after SE

To validate that UNC1999 crossed the Blood Brain Barrier (BBB) in our KA model, we performed liquid chromatography-mass spectrometry (LC/MS/MS) on plasma and hippocampal tissue from KA treated animals. UNC1999 compound was detectable in both plasma and hippocampal brain tissue after a three-day dosing period epileptogenic (**Supplemental figure 6B**).

To determine whether UNC1999 engaged its target, EZH2, to alter gene expression, we performed qRT-PCR on hippocampal tissue from saline and KA mice treated with vehicle or UNC1999. UNC1999 treated KA animals show increased gene expression in 4 out of 5 EZH2 gene targets (**Figure 7A**) and EZH2 levels were unaffected (**Figure 7B,C**). Finally, we found that 10 days after SE induction, and 8 days after the last UNC1999 dose, H3K27me3 levels are significantly reduced in the mouse hippocampus compared to no drug treatment (**Figure 7B,D**). Taken together, these results are consistent with the hypothesis that EZH2 inhibition is disease modifying and causes functional changes in gene expression and H3K27me3 levels post SE in the KA mouse model.

**Figure 7:**
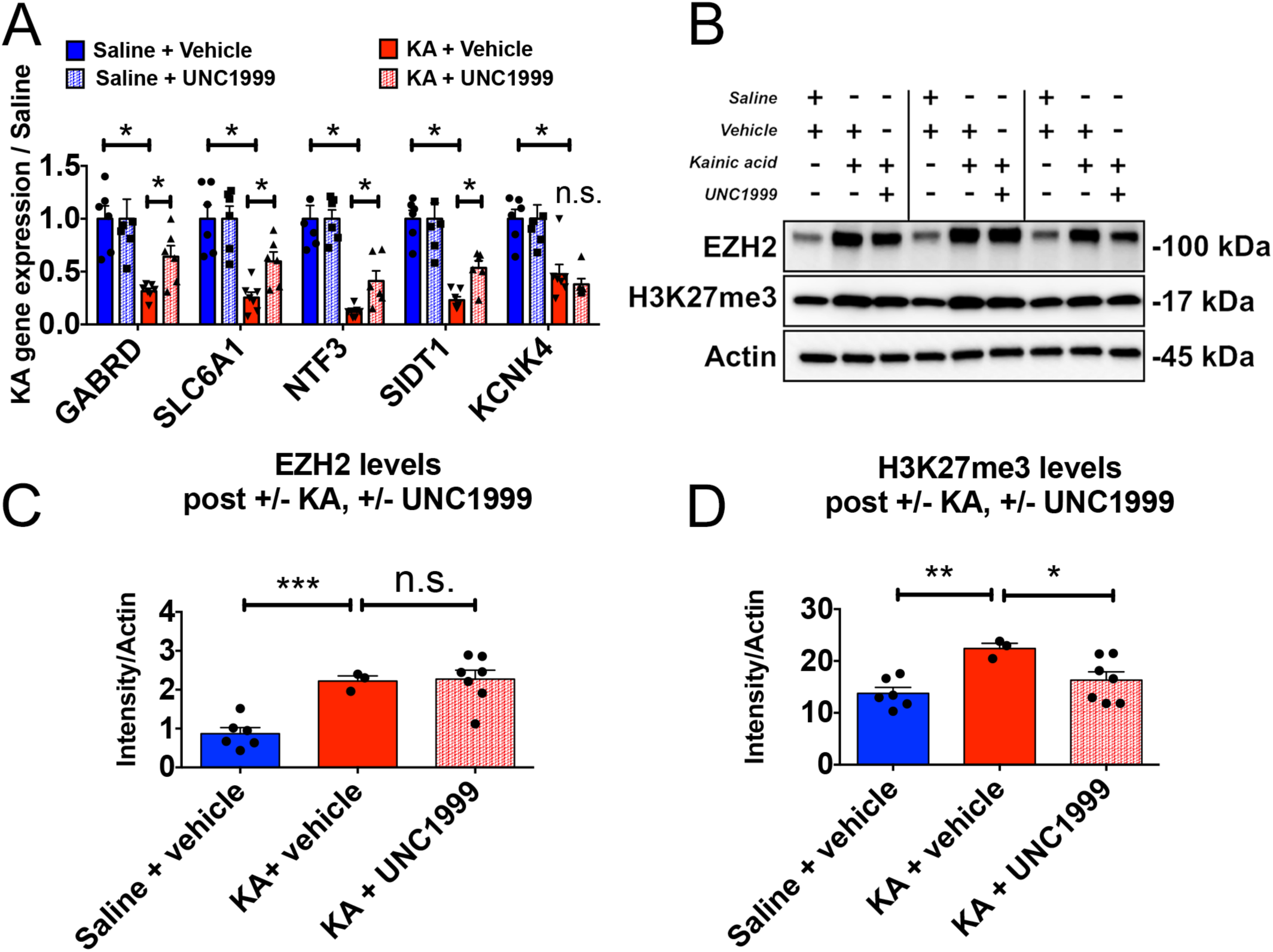
UNC1999 treatment three days *in vivo* is sufficient cause functional changes in EZH2 target gene expression and H3K27me3 levels. (A) EZH2 target genes are de-repressed after treatment with UNC1999 (multiple t-tests, p <0.05, with Holm-Sidak correction). (B) EZH2 protein levels do not change eight days after UNC1999 treatment (quantified in (C)). (D) H3K27me3 levels fail to increase at 10 days post SE in animals treated with UNC1999. Statistical significance assessed by Student’s unpaired t-test with equal SD (p < 0.05), n = 6 saline + vehicle, n = 3 KA + vehicle and n = 7 KA + UNC1999 animals. *p <0.05, **p < 0.01, ***p< 0.001, ****p <0.0001.

## Discussion

In this study, we find that EZH2 is up-regulated after SE in multiple rodent epilepsy models and provide evidence suggesting that EZH2 function is increased in human TLE. We show that transient inhibition of EZH2 acutely after SE causes an increase in seizure burden during the chronic phase suggesting that EZH2 acts as a protective agent during epileptogenesis. To our knowledge, this is the first study to characterize a functional role for EZH2 in epilepsy.

We identified EZH2 through mining a dataset provided by the Epilepsy Microarray Consortium^16^ that consists of transcriptome data from 3 epilepsy models across 11 laboratories. This consortium approach represents a novel strategy to address the problem of preclinical findings failing to translate to successful clinical trial outcomes as outlined by Landis *et al* in a 2012 Nature Perspective article^41^. By focusing on only those transcriptomic changes that occur across models and laboratories we avoided genes specific to any one experimental paradigm. Cross validation of the principle finding that EZH2 is up-regulated post SE across models (**figure 4**) demonstrates the utility of this approach.

Though it is routine to acquire whole transcriptome data, defining the transcription factors and cofactors behind large scale gene changes in an experiment remains a challenge. Motif mining near promoter regions of differentially expressed genes suffers from high false positive rates for transcription factors and does not allow for identification of cofactors^42^. Our approach of screening ChIPseq tracks archived at ENCODE^18^ for enrichment of transcription factors and cofactors at differentially expressed genes overcomes the limitations of motif mining. However, being limited to ENCODE ChIPseq tracks, we were only able to screen 161 nuclear proteins which represents about 10% of all potential human transcription factors^43^. Nevertheless, the approach identified EZH2 as a robustly induced cofactor across multiple epilepsy models and human TLE. It is a generally applicable method for defining nuclear proteins that preferentially bind, and possibly regulate, a group of genes.

The cross comparison of epilepsy transcriptomes with ChIPseq tracks archived at ENCODE allowed us to identify transcriptional regulators responsible for large-scale gene changes during the early latent period. Our analysis shows that persistently repressed genes during the early latent period are enriched for EZH2 targets. For up-regulated genes, STAT3, a nuclear factor associated with the JAK/STAT pathway, is the principal driver. This observation aligns with reports arguing that activation of the JAK/STAT pathway occurs in both pilocarpine and kainic acid SE models^44-46^. Interestingly, EZH2 was also found to be a significant driver of up-regulated genes (**Figure 2A**). This suggests that either EZH2 acts as a co-activator at these loci, or it acts to temper expression of these induced genes. We considered the possibility that EZH2 might facilitate activation of STAT3 by methylation^47,48^. However, comparison of EZH2 and STAT3 target genes do not show a significant overlap (**Supplemental Figure 4**). This suggests that either EZH2 and STAT3 do not co-localize on chromatin when EZH2 methylates STAT3, or that EZH2 does not coordinate with STAT3 in the brain as it does in cancer cells. Of note, EZH2 can act as a co-activator of other transcription factors upon phosphorylation of Serine21 in prostate cancer cells^49^ opening up the possibility that EZH2 could act as both a transcriptional repressor and co-activator in epileptogenesis.

Our analysis of Human TLE transcriptomes identified a gene module (M-1) that is enriched for EZH2 targets (**Figure 3A**). This block of 1597 genes is targeted by a group of transcription factors and cofactors that largely overlap with those that control down-regulated genes in the rodent analysis. Consistent with M-1 being under the control of EZH2, we observe a significant repression of these genes in human TLE samples compared to hippocampi of non-epileptic human samples (**Figure 3F**). This finding is remarkable given that rat transcripts were obtained in the early phases of epileptogenesis whereas the human transcripts were from patients with long-standing epilepsy. This finding raises the possibility that an epileptogenic process might continue well into the chronic phase of the disease, such that some antiepileptogenic therapies, when they are eventually identified, might also be effective in existing epilepsy.

Gene ontology analysis revealed that genes regulated by EZH2 are associated with pathways involved in neuronal function and communication, such as: “transmission across chemical synapses”, “neurotransmitter binding and downstream transmission”, “trafficking of AMPA receptors at the synapse”, and “axon guidance”. It is tempting to speculate that EZH2 upregulation in epilepsy may be an attempt to modify synaptic transmission as a protective strategy that is ultimately overwhelmed. Overall, our analysis reveals that there is a distinct gene module regulated by EZH2 in rodent and human epilepsy. Future studies will focus on assessing the role of these targets in epilepsy.

Results from the systemic KA and pilocarpine rodent models reinforce our computational predictions made from both rat and human transcriptomic data. We find that EZH2 protein levels manifest a prolonged increase in neurons but not astrocytes after SE, which peaks at 2 days and remains increased out to 5 days. EZH2 inhibition by UNC1999 significantly increases seizure burden compared to vehicle controls, suggesting a protective role for EZH2 upregulation post SE. Interestingly, EZH2 and Polycomb also play a protective role in brain ischemia. Thus, exposure to sublethal ischemic insults increases the levels of Polycomb-group proteins and overexpressing the PRC1 component BMI1 *in vitro* is sufficient to induce ischemic tolerance without requiring protective pre-conditioning^50^. These data along with findings herein may suggest a general protective role for Polycomb in brain trauma.

EZH2 missense mutations in humans cause the rare congenital disorder Weaver Syndrome, leading to neurological abnormalities such as macrocephaly, speech delay, intellectual disability, and poor coordination and balance^51^. Though not all mutations have been extensively characterized, the incorporation of some mutations into PRC2 complexes *in vitro* reduces their ability to catalyze methylation of core histones^52^. Case studies of Weaver Syndrome patients also describe the occurrence of tonic-clonic or absence seizures in adolescence with variations of both hyper- or hypo-tonia^53^. In our study, we reveal that EZH2 upregulation is neuroprotective and inhibition of EZH2 activity exacerbates disease progression. EZH2 function is a potential target for neuroprotective manipulation and disease modification during epileptogenesis.

## Methods

### Differential Expression Analysis and Clustering of Epilepsy Microarray Consortium Data

Affymetrix CEL and Rat230.cdf files were processed according to Dingledine et al, 2017^16^. Briefly, probes values were RMA normalized and gene symbols were filtered for expressed genes and those having unique probes as described in Dingledine et al, 2017^16,54^.. Of the 15,248 genes present on the array, filtering resulted in 9,614 uniquely mapped and expressed genes across the 2 pilocarpine models, 2 kainate and 1 Self Sustaining Status Epilepticus models. Each model has 6 rats per condition (control, 1-, 3- and 10-days post SE). The log2 median expression for each expressed gene in each condition was used as the expression value for that gene giving 5 values per gene for controls and 5 values per gene for epileptic rats for each day. Student t-tests were performed between controls and epileptic samples 1d, 3d and 10d post SE and p values were corrected for multiple tests^1^. Genes with FDR<0.05 and a fold change more than 2 standard deviations from the mean fold change were subjected to hierarchical clustering using uncentered Pearson correlations and complete clustering.

### Identification of Transcription Factors and Cofactors

A single ChIP value was assigned to each Factor in ENCODE as follows: All genes in the human genome (hg19) were assigned a gene domain that was defined as 5Kb either side of the gene body (5’UTR – 5Kb to 3’UTR + 5Kb). NarrowPeak files for all transcription factors and cofactors were downloaded from 2012 data freeze (last updated 21-July-2013) http://genome.ucsc.edu/cgi-bin/hgFileUi?db=hg19&g=wgEncodeAwgTfbsUniform and custom Python scripts were used to extract the highest ChIPseq peak value (signalValue) for a given factor seen in any cell line within every gene domain. Thus, each factor is assigned a single signalValue per gene – the highest value – observed in ENCODE. The resulting gene x factor array was normalized such that the sum of all signalValues for any factor was arbitrarily set at 100,000.

### Defining Enriched Factors in a Gene List

For each Factor in the above array, the 9614 expressed genes (background list) are ordered by ChIP signal. A normalized cumulative distribution function is generated that accumulates the number of genes with increasing ChIP signal from zero to the maximum signal in the array for that Factor:

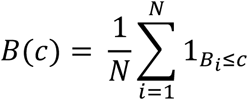

Where B(c) is the fraction of genes in the background list (N genes) with ChIP signal less than or equal to c. B are the ordered ChIP signals.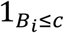 is an indicator function that is equal to 1 if B_i_ ≤ c else equal to 0. An emperical distribution function, Q(c) is generated by accumulating the ordered X query genes (Q) (where Q ⊂ B and 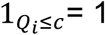 if Q_i_ less than or equal to c else equal to 0):

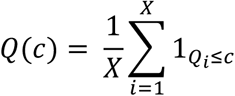

We then screen for Factors where the query CDF is right-shifted compared to the background CDF. This is accomplished by calculating the difference between the 2 distributions and defining the supremum (D_S_) and infimum (D_I_) of the difference as:

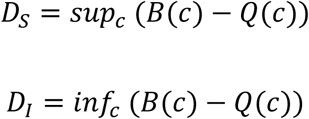

If the cumulative of query samples is left shifted compared to the population (|D_S_| < |D_I_|), the factor is triaged and not considered further. If the query cumulative is right shifted compared to the population cumulative i.e. |D_S_| > |D_I_|, a 1-tailed Mann-Whitney-Wilcoxon (MWW) test between the background and query list is performed.

A Score for each Factor is calculated to incorporate the Benjamini-Hochberg corrected MWW p value, as well as a measure of how the highest ChIP values in the query list compare to the highest values in the background list. Thus, for each Factor, the mean of the top n chip signals in the query list of length X, is compared to the mean of the top n ChIPs in the population to produce a ratio such that:

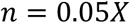

and

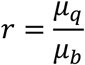

Where μ_q_ is the mean for the top n signals for Q and μ_b_ is the mean of the top n signals for B. μ_q_ and μ_b_ are referred to ‘Obs Tail Mean’ and ‘Exp Tail Mean’ in the supplemental factor analysis xls files.

A Score, S, is assigned to each Factor thus:

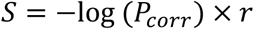

Where P_corr_ is the Benjamini-Hochberg corrected MWW 1-tailed p value. Factors are then sorted by Score.

### Analysis of overlap of target genes for each Factor

A subset of the input list is preferentially targeted by a given Factor i.e has higher ChIP signals than the whole input list overall. Target genes are defined as all genes with ChIP signals greater than the argument of d_sup_ as defined by the blue vertical blue lines in **Supplementary Figure 2.** This is equivalent to the argument of the Kolmogorov-Smirnov statistic for the 2 curves and analogous to a ‘Leading Edge’ analysis in GSEA^23,24^. The argument of d_sup_ is refered to as the ‘Critical Chip’ value in the supplemental factor analysis xls files. To determine the degree of overlap between target genes for each positive Factor, Fisher Exact tests (Venn analyses) are performed for all pairwise combinations of target genes to assess overlap between lists. Those Factors with significant overlap of target lists (odds ratio>1 and p<0.05) are displayed on the heatmaps in Figure 2 as shades of red. Factors with odds ratio ≤1 or p≥0.05, are designated white on the map. The resulting array is then clustered by Pearson distance and Factors are considered as grouped if their Pearson distance is less than 0.7 (Pearson correlation>0.3).

### Analysis of human Temporal Lobe Epilepsy and control datasets

#### Factor analysis of human TLE samples

Dataset GSE63808^15^ containing 129 ante-mortem transcriptomes from TLE resected tissue was downloaded from GEO. Probes with an associated Illumina p-value<0.05 in at least 20% of samples were kept for further analysis. Probes were collapsed to gene symbols by taking the median probe expression value per symbol. Expression values were column-centered and normalized by dividing each value in a sample by the median expression value in that sample and weighting each column equally. Genes with variance across samples greater than the median coefficient of variance across samples were taken for further analysis resulting in 7658 unique gene symbols. K-medians analysis was used to divide genes into 10 clusters (M-1 to M-10). Each cluster was used as an input gene list for Factor analysis with the 7658 unique genes as background.

#### Comparison of human TLE k-medians clusters and rodent up- and down-regulated gene sets

Factor analysis FDRs for the k-medians human TLE clusters and the 6 rat gene lists (up and down genes at 1d, 3d and 10d) were combined into a 2-dimensional array (columns (x) = gene lists, rows (y) = Factors). Thus FDR_**x,y**_= Factor analysis FDR for gene list **x** and Factor **y**. The array was digitized such that the array value for gene list **x** and Factor **y** (V_**x,y**_)= 1 (black) if FDR_**x,y**_ < 0.05 else V_**x,y**_ = 0 (white). Columns were then clustered using Pearson correlation.

#### Analysis of non-epilepsy human hippocampus transcriptomes

GSE45642 was downloaded from GEO^30^. Samples 1-55 (hippocampal control samples with no psychiatric or neurological disorders, substance abuse, or any first-degree relative with a psychiatric disorder) were kept for analysis. Expression values were column-normalized by dividing each value in a sample by the median expression value in that sample. M-1 genes were extracted and the average expression of each gene in M-1 across samples 1-55 was compared to the average expression of each gene in M-1 across the TLE samples using a Mann-Whitney test.

#### Animal care

All animal procedures were performed with approval from the University of Wisconsin-Madison School of Medicine and Public Health Institutional Animal Care and Use Committee and according to NIH national guidelines and policies.

#### Low-dose KA mouse model

Male FVB/NHsd mice (6 weeks old) were ordered from Envigo (Madison, WI) and housed under a 12-hour light/dark cycle with access to food and water *ad libitum*. Animals were allowed to recover for at least 48 hours after transport before engaging in experimentation. All KA experiments were performed at the same time of the day (∼9 A.M.) and mice were returned to their home cages by ∼6 P.M. All mice were singly housed upon arrival and after induction of SE due to aggressive behavior.

We utilized a low-dose KA mouse model to induce SE. To begin, mice were first intraperitoneally (ip) injected with 10 mg/kg of KA diluted in Milli-Q water or 0.9% saline (Tocris Bioscience, Bristol, United Kingdom). After twenty minutes, mice were given a second dose of KA at 5 mg/kg. Animals continued to receive 5mg/kg doses of KA every twenty minutes until each reached SE. During the induction process, epileptic behavior was scored using a modified version of the Racine Scale^40^, where 1 = freezing, behavioral arrest, and staring spells; 2 = head nodding and facial twitches; 3 = forelimb clonus, whole body jerks or twitches; 4 = rearing; 5 = rearing and falling; 6 = continuous rearing and falling; and 7 = violent running or jumping behavior. Animals were declared to be in SE when at least five Racine 5-7 level seizures were observed in a 90-minute time window. KA mice were maintained in SE for three hours and all animals, both KA and saline treated, were later IP injected with 5 mg/kg of Diazepam (Hospira Inc., Lake Forest, IL) to relieve seizure burden and reduce mortality. To ensure proper recovery, animal cages were placed on a heating blanket after diazepam treatment for 1-2 hours. After recovery, all animals were IP injected with 0.5 mL of 0.9% saline and soft gel food was provided in home cages.

#### Lithium-pilocarpine rat model

Seizures were induced in adult Sprague Dawley rats (8 weeks old, 250-300 g) from Envigo. To induce status epilepticus, rats were injected with lithium-chloride (127 mg/kg, IP, Sigma, St. Louis, MO) 24 hours prior to pilocarpine hydrochloride (50 mg/kg, IP, Sigma). Methylscopolamine (1 mg/kg, IP, Sigma) was administered 30 minutes prior to pilocarpine to reduce peripheral cholinergic effects. Rats that did not exhibit behavioral seizures within 1 hour of pilocarpine injection were given a second dose of 25 mg/kg pilocarpine hydrochloride. Motor seizures were scored by standard behavioral classes as follows: (1) behavioral arrest, eye closure, vibrissae twitching, sniffing; (2) facial clonus and head bobbing; (3) forelimb clonus; (4) rearing with continued forelimb clonus; and (5) rearing with loss of motor control and falling. Rats were treated subcutaneously with 6 mg/kg Diazepam after one hour of seizures.

#### Hippocampal tissue isolation and homogenization

After KA seizure induction, animals were sacrificed, whole hippocampi hemispheres were harvested, and flash frozen in liquid nitrogen. A single hippocampal hemisphere was lysed in Radioimmunoprecipitation Assay buffer (RIPA: 50mM Tris, 150mM NaCl, 1% nonidet P-40, 0.5% sodium deoxycholate, 0.1% SDS) with mammalian protease inhibitor (1:100, Sigma) and phosphatase inhibitors (10mM NaF, 2mM Na Orthovanadate, 4mM Na pyrophosphate, 10mM Na β-glycerophosphate). Tissue was homogenized using a probe sonicator (Fisher Scientific, Sonic Dismembrator, Model 100, Hampton, NH) by sonicating on power 4 for three rounds with 10 pulses each round. Samples were then centrifuged at 13.5 x g for 30 minutes to separate cell debris. Supernatants were isolated and quantified using the DC Protein Assay (Bio-Rad, Hercules, CA). 5X loading buffer (0.5M Tris, 10% SDS, 50% glycerol, 10mM EDTA and 1% B-mercaptoethanol) was added to each sample to reach a 1X final concentration. Protein extracts were boiled at 95°C for 5 minutes and stored at −80°C until run on an SDS PAGE gel.

#### Western Blotting

Protein extracts were loaded at 20μg per lane and resolved by standard electrophoresis in 4-20% Mini-PROTEAN TGX Precast polyacrylamide protein gels (Bio-Rad, Hercules, CA). Gels were transferred onto polyvinyl difluoride membranes (PVDF; Millipore, Bedford, MA) using Tris-glycine transfer buffer (20mM Tris, 1.5M glycine, 20% methanol). Membranes were blocked with 5% non-fat dry milk diluted in low-salt Tris-buffered salt solution (w/w TBST; 20mM Tris pH 7.6, 150mM NaCl, 0.1% Tween 20) for 1 hour at room temperature. Primary antibodies were diluted in 5% non-fat dry milk in TBST and incubated with membranes overnight at 4C. Antibodies include: EZH2 (1:1000, Cell Signaling, Danvers, MA), EED (1:1000, Active Motif, Carlsbad, CA), SuZ12 (1:1000, Cell Signaling), H3K27me3 (1:1000, Active Motif), and Actin (1:10,000, MP Biomedicals). The next day, membranes were washed three times with 1X TBST and incubated with horseradish peroxidase-conjugated goat anti-rabbit or goat anti-mouse IgG secondary antibodies (1:10,000, Santa Cruz Biotech, Dallas, TX) for 1 hour at room temperature. Afterwards, membranes were washed three times with TBST, and protein bands were detected using SuperSignal West Femto ECL reagent (ThermoFisher, Waltham, MA). Bands were visualized using a ChemiDoc-It Imaging System (UVP VisionWorks, Upland, CA) and quantified using UVP Vision Works software. Band intensity quantifications were graphed and analyzed using GraphPad Prism (GraphPad Software, La Jolla, CA).

#### RNA extraction and Quantitative Real-time PCR (q-RTPCR)

RNA from flash frozen hippocampal tissue was extracted using TRIzol reagent (ThermoFisher, Waltham, MA) according to the manufacturer’s protocol, resuspended in 20ul of sodium citrate (1mM, pH 6.4) with 1X RNA secure (ThermoFisher) and quantified using a NanoDrop Spectrophotometer (ThermoFisher). 1ug of isolated RNA was reverse-transcribed using SuperScript III (Invitrogen, Carlsbad, CA) according to manufacturer’s instructions. Each reverse transcription reaction was diluted 1:4 with molecular grade water. Quantitative real-time PCR was carried out using SYBR Premix Ex Taq (Takara Bio Incorporated, Kusatsu, Shiga Prefecture, Japan). Primers used for qRT-PCR were as follows: **EZH2**, forward 5’ GGC TAA TTG GGA CCA AAA CA 3’ and reverse 3’ GAG CCG TCC TTT TTC AGT TG 5’, **SLC6A1** forward 5’ CAT TGT GGC GGG CGT GTT 3’ and reverse 3’ CTC AGG GCG CAC AAT ATC 5’, **GABRD** forward 5’ CGC CTA CAG CCT GAT GGG GTG ATT 3’ and reverse 3’ GGG AAC TGG CCA GCC GAT TTG AAG 5’, **SIDT1** forward 5’ TCG CCA GCA GAA AGA AGT 3’ and reverse 3’ TGA GAG GGG CTG GCA GTG 5’, **NTF3** forward 5’ CAC GGA TGC CAT GGT TAC TTC TGC 3’ and reverse 3’ GTG GCC TCT CCC TGC TCT GGT TC 5’, and **KCNK4** forward 5’ CCG GGG CTG GTG AGA AGT 3’ and reverse 3’ GCG GCT GGT AGG CTG GAG 5’.

#### Perfusion and sectioning of KA and saline mouse brains

KA and saline mice were sedated by vaporizing isoflurane and then intra-cardially perfused with 1X PBS followed by 4% paraformaldehyde (PFA; Electron Microscopy Sciences, Hatfield, PA). four days post SE (n = 3 KA and n = 3 saline mice per condition). Toe and tail pinch were utilized to ensure complete and proper anesthesia. A 22-gauge butterfly needle (Becton Dickinson, Franklin Lakes, NJ) was pushed into the apex of the heart and the right atrium was slashed upon perfusion of 1X PBS. Whole brain dissection was performed, and brains were fixed for an additional 24 hours in 4% PFA at 4°C. Brains were embedded in 6% agarose and sectioned coronally at 30-micron thickness using a Leica VT1000S vibratome (Leica Camera, Wetzlar, Germany).

#### Immunofluorescence staining and confocal microscopy

Free-floating hippocampal sections were treated for an antigen retrieval process. Briefly, sections were incubated in 10 mM Sodium Citrate Buffer (pH 8.5) and at 80°C for 30 minutes. Next, sections were blocked in NGS blocking solution (10% Normal Goat Serum, 0.4% Triton X-100, 1% Glycine, 2% BSA in Tris-Buffered Saline [1M Tris pH 7.5, 5M NaCl]) at room temperature for 2 hours with gentle agitation. Primary antibodies EZH2 (Cell Signaling, 1:200), NeuN (Abcam, 1:500) or GFAP (Abcam, 1:500) were added directly to wells and incubated overnight at 4°C with gentle agitation.

The following day, sections were washed three times with 1X PBS and incubated with secondary antibodies goat anti-rabbit AlexaFluor488 (Invitrogen, 1:500) and donkey anti-mouse AlexaFluor594 (Invitrogen, 1:500) diluted in blocking solution for 3 hours at room temperature. Sections were washed with three times with 1X PBS and counterstained with DAPI (Thermofisher, 1:500 from 5mg/mL stock) at room temperature for 30 minutes. Sections were washed twice with 1X PBS, mounted onto gelatin-coated slides (Southern Biotech, Birmingham, AL) and cover slipped using Prolong Gold Mounting Media (Invitrogen). Slides were imaged at 60x oil magnification using a Nikon A1Rs confocal microscope (Nikon, Tokyo, Japan). Images were analyzed using the NIS elements program.

#### UNC1999 treatment of KA mice

UNC1999 (Cayman Chemical, Ann Arbor, MI) was diluted at 2.5 mg/mL in 4% N, N-Dimethylacetamide (DMA; Sigma), 5% Solutol (Sigma) and 91% normal saline drug vehicle. To dissolve UNC1999, DMA was added to UNC1999 and vortexed to mix. Once dissolved, UNC1999 was added to a 0.9% saline solution containing Solutol and bath sonicated twice for twenty-minutes each at 65°C. Mice were treated with 20 mg/kg UNC1999 or drug vehicle six hours (3 hours post diazepam), 24 hours, and 48 hours post SE.

#### Video recording of epileptic behaviors in KA mice after vehicle and UNC1999 treatment

After a latent period of 4 weeks, KA + UNC1999 and KA + vehicle treated cohorts were placed in housing units containing nine cubicles. Housing units were organized in a vertically stacked, 3×3 unit fashion, with transparent plastic walls on either side of each unit, and opaque floors. Each cubicle included bedding and free access to water and food *ab libitum*. To quantify instances of epileptic seizures, mice were video recorded from the hours of 9 A.M. to 5 P.M., 6 days a week for 3 weeks. Each treatment group was assigned one camera, which was set in front of each housing unit. Video recording began automatically at 9 A.M. each morning.

Two blinded observers (one male, one female) used a modified version of the Racine Scale to score and quantify instances of behavioral seizures. Experimenters decoded and unblinded observers after completing all video scoring and analysis.

#### Statistical Analysis

For all tests, p < 0.05 adjusted for multiple comparisons was considered statistically significant. All data were graphed as the mean ± standard error of the mean (SEM) in Prism 6 software (Graphpad Software, La Jolla, CA). Statistical tests were also performed using Prism 6 software. Western blot, RTPCR and video recording data were subjected to 2 way ANOVA with Holm-Sidak correction for multiple comparisons.

#### Western Blots

Protein bands from Western Blots were quantified using the UVP Vision Works software. EZH2, SUZ12, EED, and Total H3 quantifications were all normalized to Actin quantifications. H3K27me3 quantifications were normalized to total H3 quantifications.

#### Quantitative PCR for mRNA expression levels

Gene expression levels were calculated for every sample using the Delta Ct method and normalized to B-actin expression levels. Data were graphed by taking the average of the normalized expression levels of all saline samples, and then dividing each sample value by that average number to set control levels at unity.

#### Video Recording

Behavioral seizures were scored by blinded observers and totaled for each individual animal at each Racine stage displayed every day of the recording. These numbers were utilized to analyze the outcome of drug treatment in Prism 6 software. The average number of daily seizures exhibited by each treatment group was calculated by adding the total number of scored events exhibited by all (n = 8) animals in each treatment group every day of the recording. Students unpaired t-test with equal SD was applied to assess significance. There was a total of 18 days of recording across 3 weeks. Recordings were performed six out of the seven days of each week. Bedding, water, and food was maintained on day 7. The number of seizures exhibited by each treatment group per week was calculated by adding up the total number of scored events exhibited by all n = 8 animals in each treatment group.

The number of events exhibited in each Racine stage for each treatment group was calculated by adding up all R1-R7 events scored for each animal during the entire three weeks of recording.

#### LC/MS/MS

Concentrations of UNC1999 detected in plasma and brain were expressed as ng/mL after LC/MS/MS. To determine how many nanograms of drug was detected per gram of hippocampal tissue, all hippocampal samples were weighed prior to homogenization for LC/MS/MS and the average was noted. Concentration of drug detected was divided by the average brain tissue weight and expressed as ng of drug/g wet weight of tissue. Values were plotted on a log base 10 scale. Plasma values were expressed as ng/mL after LC/MS/MS and were plotted on a log base 10 scale.

## Figures

**Supplemental figure 1:**
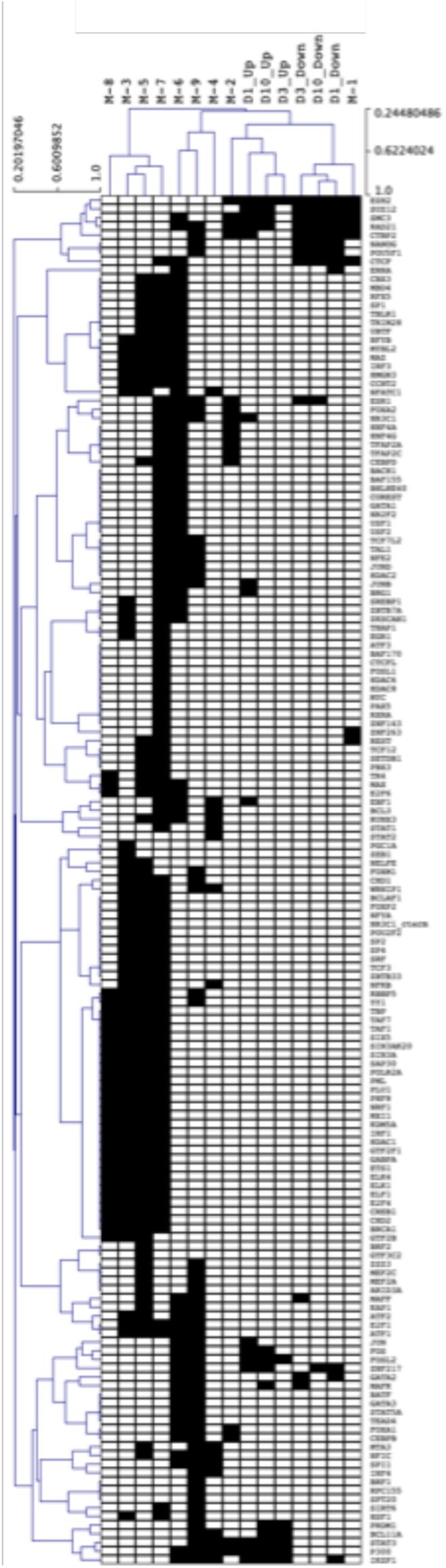
Clustering of human gene modules with gene lists up- or down-regulated at 1,3 and 10 days post SE. All 10 k-median modules and gene lists for up or down regulated genes on days 1,3 and 10 were clustered based on absence or presence of factors in their respective Factor analyses. Black box denotes Factor with FDR<0.05, white denotes Factor with FDR≥0.05.

**Supplemental figure 2:**
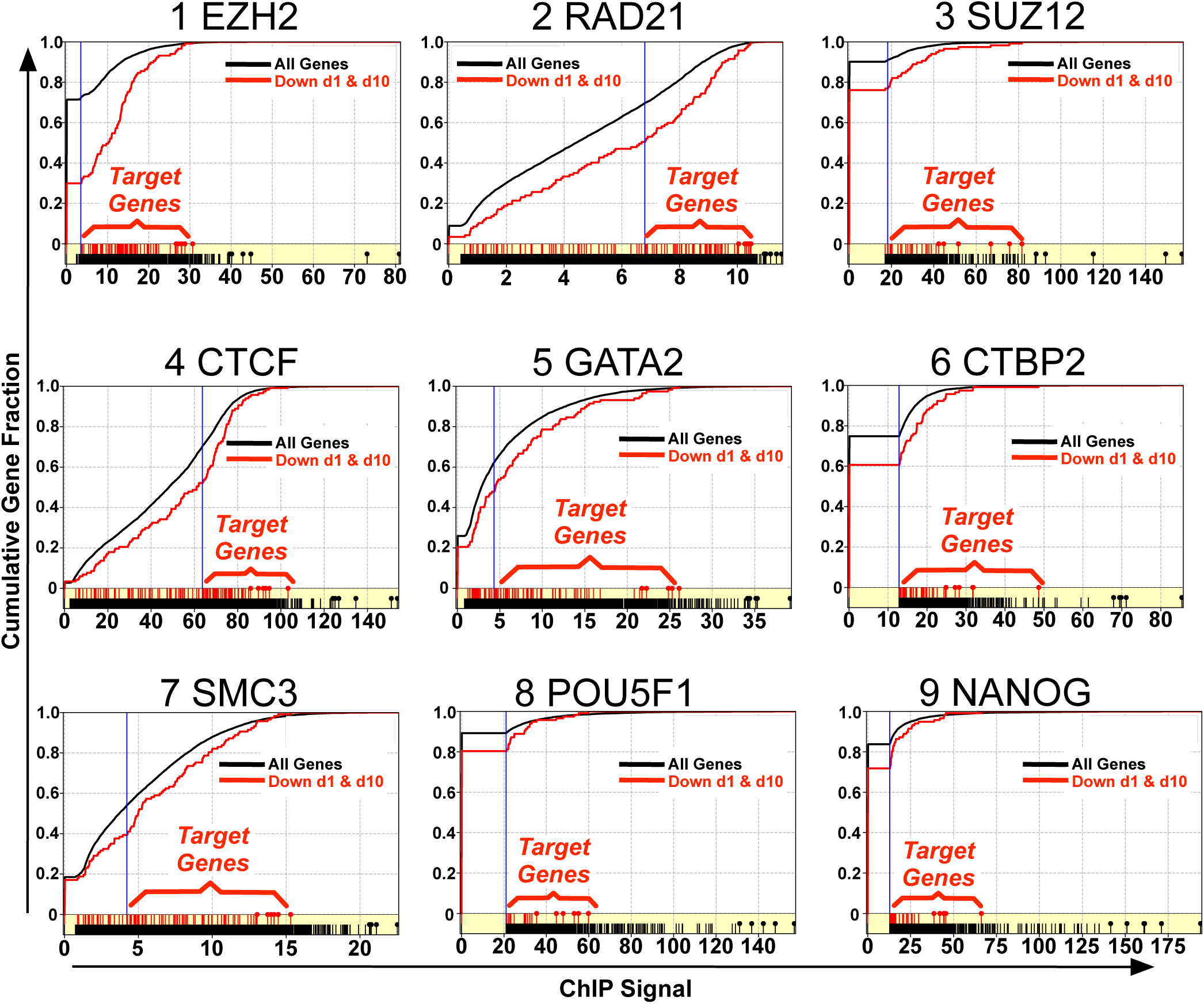
Target gene definition for Factors targeting repressed genes. Cumulative Distribution Functions of background and repressed gene ChIP signals for Factors driving gene repression on days 1 and 10 post SE. Two cumulative functions are displayed: the black curve is the fractional cumulative of all genes in the background list against ChIP values. Red is the same for fractional cumulative of all genes repressed 1 and 10 days post SE against ChIP values. A blue vertical line denotes the ChIP value at d_sup_ i.e the argument of the Kolmogorov-Smirnov statistic, representing the ChIP signal at which the largest separation occurs between the two cumulative density plots. Red ticks represent each gene in the query list and black ticks are all genes in the population. Red ticks with circles represent those genes repressed on days 1 and 10 with the top 5% ChIP signals for the Factor. Black lollipops are the top 5% genes in the Background list.

**Supplemental figure 3:**
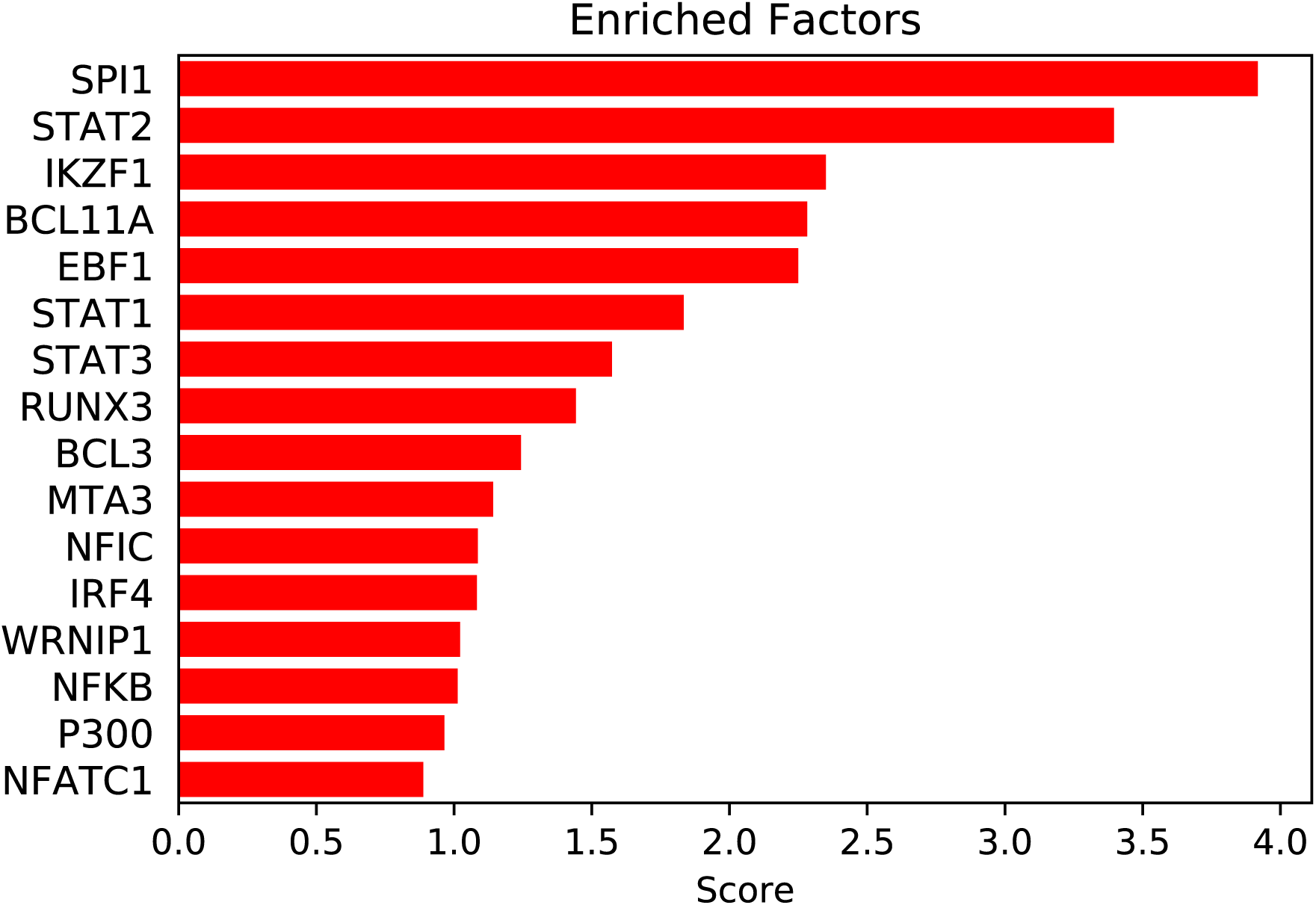
Factor analysis output for gene module M-4 in human TLE samples highlights transcription factors and cofactors with known roles in inflammation.

**Supplemental figure 4:**
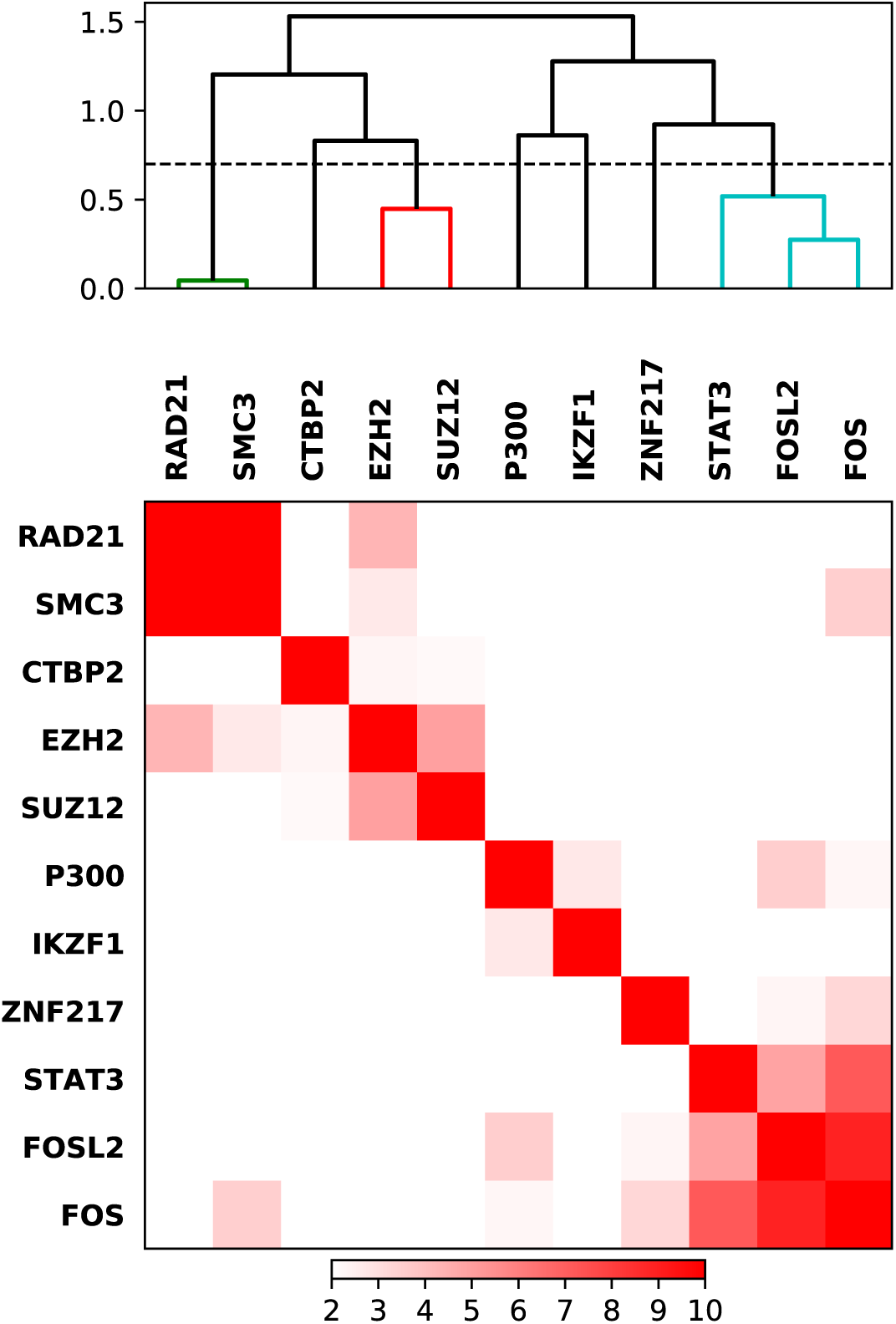
Overlaps between lists of target genes for each significant Factor driving persistently up-regulated genes are plotted as a heatmap based by Fisher Exact test -log(p) values. Clustering by Pearson distance highlights 3 groups of factors controlling persistently repressed genes. Dashed line cutting branches of the hierarchy tree at Pearson distance=0.7 (Pearson correlation=0.3).

**Supplemental Figure 5:**
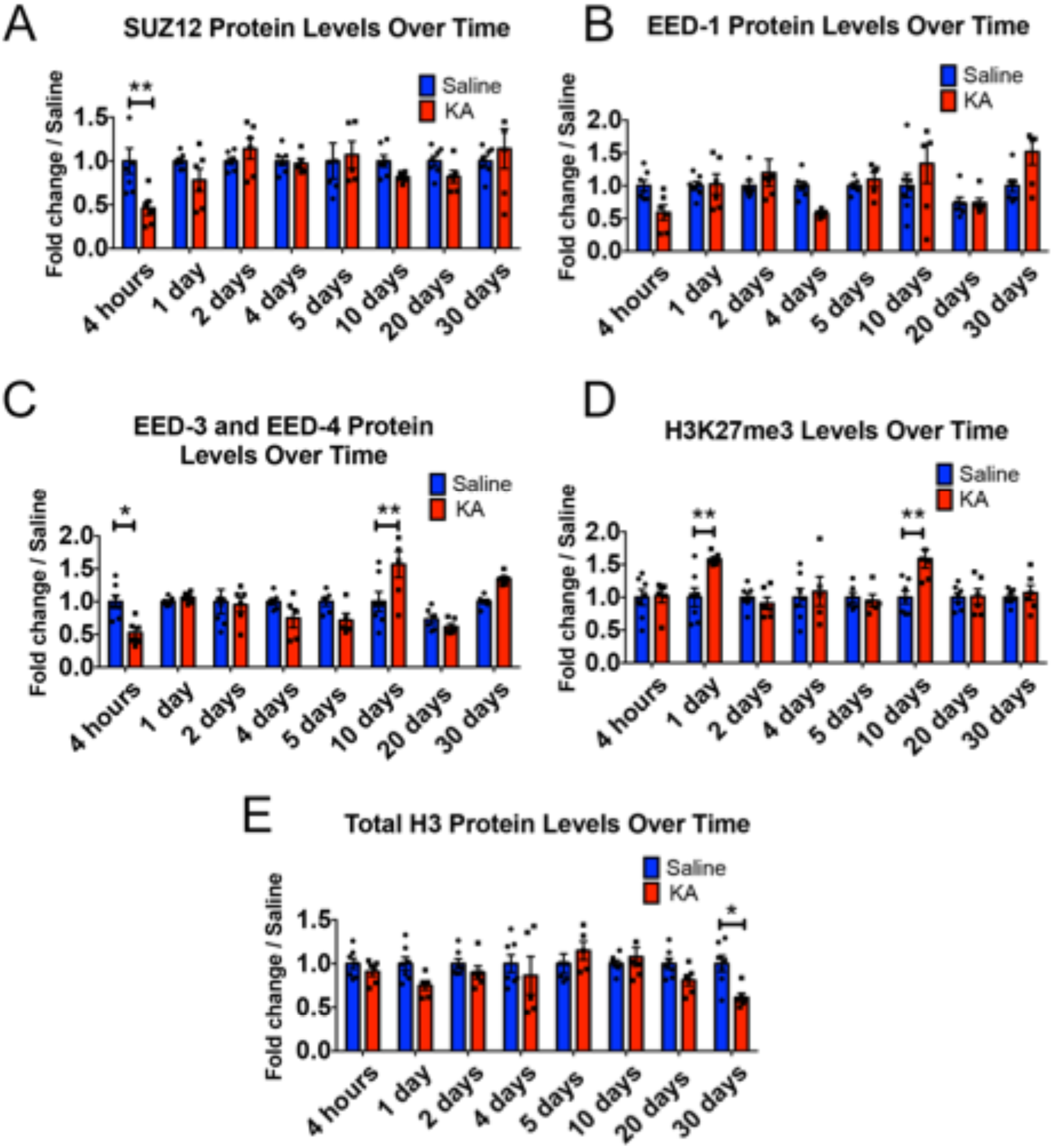
Quantification of SUZ12, EED-1, EED-3, EED-4, H3K27me3 and Total H3 western blots. (A) SUZ12 demonstrates an early, transient decrease in protein levels 4 hours after SE and remains unchanged at remaining time points compared to saline controls. (B) EED-1 is not significantly changed at any of the above tested time points. (C) EED-3 and EED-4 exhibit an early, transient decrease in protein levels 4 hours after SE, and demonstrate an increase later at 10 days. (D) The characteristic histone modification mark of EZH2 activity, H3K27me3, is significantly increased at 1- and 10-days post SE. (E) Total Histone 3 protein levels remain unchanged immediately after SE. A significant decrease in total H3 protein was detected at 30 days. Statistical significance was assessed across animals, time, and conditions through Two-way ANOVA test where p < 0.05 with Holm-Sidak correction for multiple comparisons. n = 7 for saline treated animals and n = 5-7 KA animals at every time point.

**Supplemental Figure 6:**
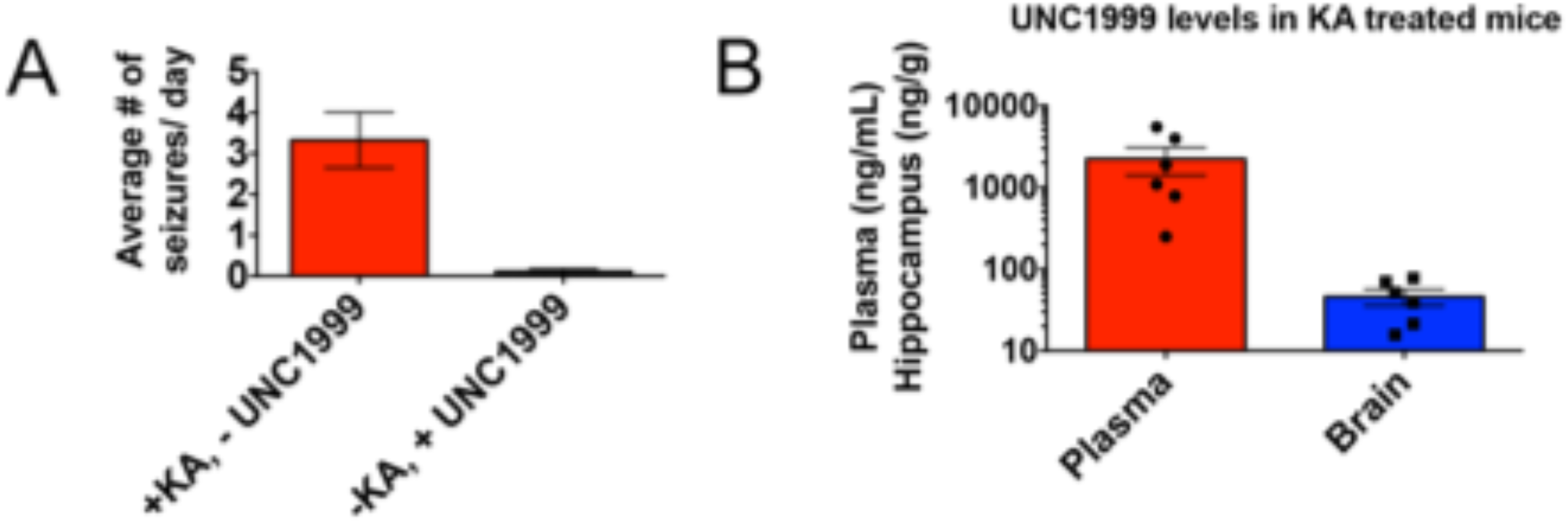
(A) UNC1999 is not a frank convulsant to naive animals that are treated with UNC1999 only (i.e. no KA) and then assessed for behavioral seizures by video recording. Scores were assessed by the same set of blinded observers. (B) LC/MS data from plasma and hippocampal samples from KA + UNC1999 treated mice demonstrating UNC1999 was detectable in the brain post injection.

## Acknowledgements

We would like to acknowledge Anqi Ma for providing his advice on UNC1999 drug vehicle formulations for intraperitoneal injections. We would also like to acknowledge the UW Biotechnology Center for their assistance in performing LC/MS/MS. During the course of this study, NK was supported by NIH National Research Service Award T32 and the NIH NINDS Blueprint Diversity Specialized Pre-Doctoral Award F99. AR and RD were supported by NIH R21NS093364. AR was supported by a CURE Challenge award and a Spark award from Lily’s Fund (https://lilysfund.org).

